# Urban greenspace aerobiomes are shaped by soil conditions and land cover type

**DOI:** 10.1101/2024.01.12.575340

**Authors:** Joel E. Brame, Craig Liddicoat, Catherine A. Abbott, Christian Cando-Dumancela, Jake M. Robinson, Martin F. Breed

## Abstract

Growing evidence suggests that exposure to microbial biodiversity is important for human immunoregulation and health. Urban greenspaces harbour airborne bacterial communities (aerobiomes) with the potential to transfer beneficial bacteria to humans. However, limited studies have examined the ecological influences of soil, vegetation, and rainfall on aerobiomes in urban greenspaces. Here, we utilised 16S rRNA amplicon sequence data to analyse the effects of land cover, soil abiotic characteristics, surrounding vegetation diversity, and rainfall on aerobiome diversity and composition from 33 urban greenspace sites in Adelaide, South Australia. We sampled air and soil from two urban greenspace land cover types: highly-managed sports fields (*n* = 11) and minimally-managed nature parks (*n* = 22). Sports field aerobiomes had a distinct aerobiome community composition and higher alpha diversity than nature parks. Aerobiome alpha diversity was shaped more by soil abiotic characteristics, particularly soil pH and iron levels, than woody plant species diversity. Rainfall prior to sampling also had strong effects on the aerobiome community composition and associated with decreased alpha diversity. These findings point toward soil iron and pH management as pathways to increase aerobiome bacterial diversity. Our study shows that, with additional research, there is potential for greenspace managers and urban planners to target specific soil abiotic characteristics in urban greenspaces to improve microbiome-mediated urban health.

## INTRODUCTION

Biodiverse microbial exposure via the outdoor environment is interconnected with human health through historical and co-evolutionary mechanisms, including immunoregulation [1–3]. However, the rapid expansion of modern urbanisation is leading to a consequential change in everyday environments [4]. For city residents, urban greenspaces maintain opportunities for exposure to biodiversity [5] and have been associated with improvement in physical, social, and psychological health [6]. There are various pathways linking biodiversity to human health. These include social (e.g., greenspaces provide areas for convivial activities), biological (e.g., exposure to stimuli that trigger physiological effects), psychological (e.g., exposure to stimuli that change moods and emotions), physical activity (facilitating exercise), and environmental buffering pathways (e.g., trees providing cooling that reduces heat stress; [7]). The biological pathway linking biodiversity to health is thought to encompass exposure to greater microbial biodiversity and specific microbial taxa (also known as “old friends”) that regulate the innate immune system, as described by the biodiversity hypothesis [8]. Growing evidence links greater environmental microbial diversity exposures with increases in immunomodulatory cells, including regulatory T-cells, which provide protection from allergies and some chronic inflammatory diseases [9, 10]. Thus, evaluations of urban greenspace soils, vegetation, and air for microbial diversity and composition, alongside the analyses of pathogens, can provide insights into their potential linkages with human health.

Airborne bacterial community (aerobiome) biodiversity has been examined across greenspace land covers, but the number of such studies is limited. The alpha diversity of aerobiomes has been compared across several urban land cover types, including parks versus parking lots [11], grassy versus forested areas [12], and bare ground versus grasslands and scrub habitat [13]. Dispersal from leaf surfaces is recognised as an important factor in shaping aerobiome communities [14]. It is well-established that vegetation composition directly impacts – and is impacted by – microbial communities in outdoor urban ecosystems, especially within soils [15]. Urban greenspace vegetation can take many forms; however, two contrasting and commonly encountered land cover types include: intensely maintained grass sports fields, and minimally maintained and more natural vegetation systems such as parklands or woodlands. Despite the regular human usage of sports fields and the associated potential microbial exposure, to our knowledge, they have received minimal attention in aerobiome studies.

Furthermore, the ecological influences on aerobiome biodiversity extend beyond vegetation and land cover. Many ecological variables influence the urban aerobiome, including anthropogenic activity [16], wind-carried bacterial contributions from allochthonous (distant) sites [17], and soil physicochemical conditions [18]. Temporal variation [19], vertical microbial stratification dynamics [20], and air pollution [21], among other ecological and meteorological variables [14], also modulate aerobiome composition. Analysis of these complex ecological interactions from the perspective of beneficial bacterial exposure has only recently gained momentum, and significant knowledge gaps remain. Soil abiotic/physicochemical characteristics (e.g., pH, nutrients) are known to shape the composition and stochastic and deterministic assembly of soil microbial communities [22]. However, their association with aerobiome composition has received limited attention.

Here, we characterised aerobiome community profiles along with vegetation and soil physicochemical parameters from 33 urban greenspaces across metropolitan Adelaide, South Australia, to determine the influence of local plant and soil conditions in shaping aerobiomes. Our urban greenspaces were grouped into two types: highly-maintained grassy sports fields (*n* = 11) and minimally-maintained nature parks (*n* = 22). To understand ecological influences on aerobiomes in urban greenspaces, we aimed to evaluate the effects of land cover type, woody plant species diversity, rainfall, and soil physicochemical parameters on **(1)** the aerobiome alpha diversity and **(2)** the aerobiome community composition, including specific commensal taxa.

## MATERIALS AND METHODS

### Study locations

We sampled spatially independent 25 x 25 m replicates of sports fields (*n* = 11) and nature parks (*n* = 22) (33 sites total) in greater metropolitan Adelaide, South Australia (**Fig. 1a**). The 25 x 25 m area is shown to be suitable to describe vegetation and microbial variation [15, 23]. A panoramic photo was taken at the centre of each replicate (**Fig. 1b-1c** shows examples of greenspace types). Sites were chosen so that: (1) all sites were at a distance of > 5 km from the coast to avoid coastal effects; (2) all sites were within the low-elevation plains metropolitan Adelaide region to avoid the different (colder, wetter) climatic conditions of the hills and mountains bordering to the east; and (3) nature park sites represented a range of woody plant diversity.

**Fig. 1.**
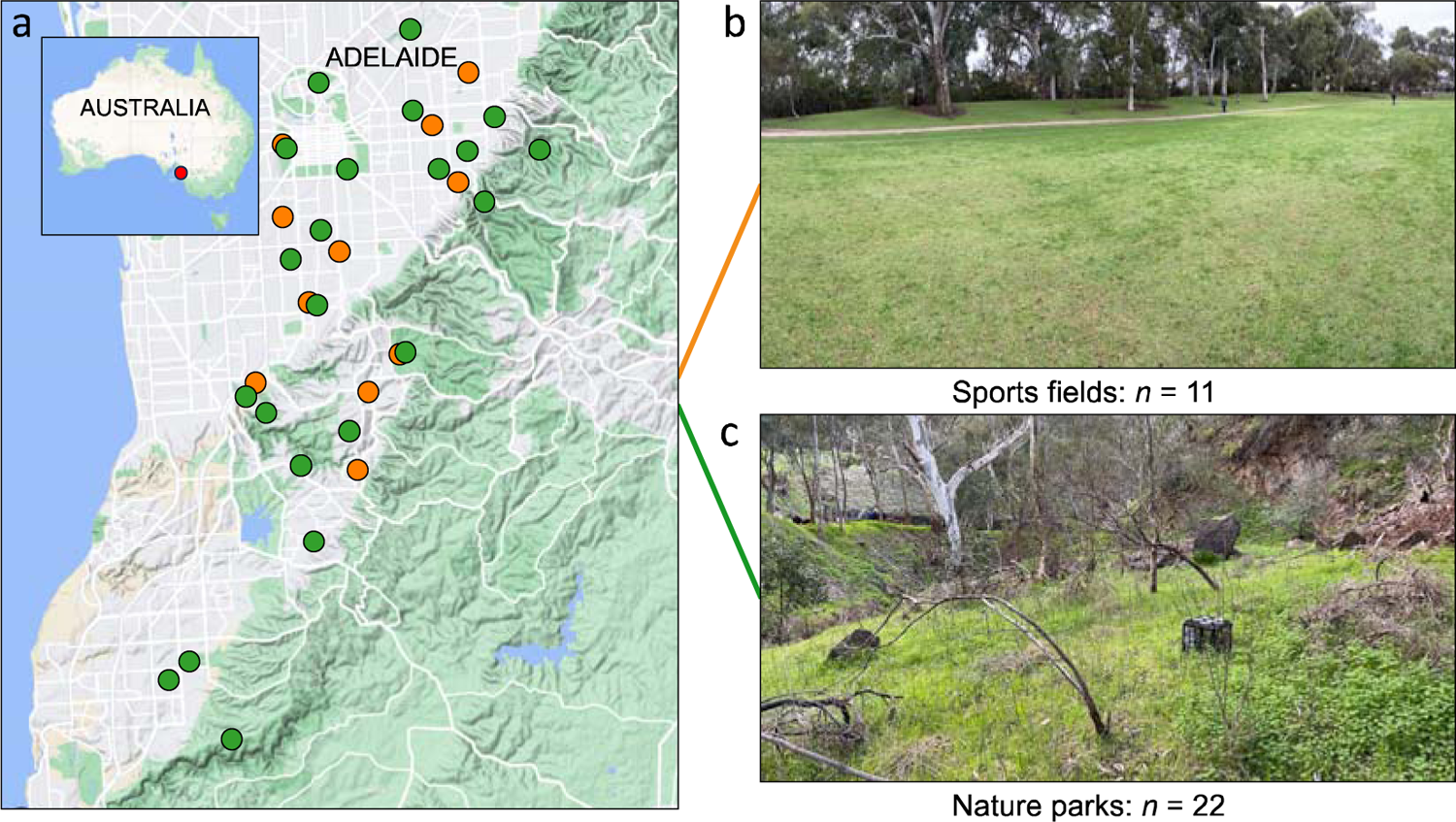
(a) Map of Australia and sampling sites across greater metropolitan Adelaide. Orange dots denote sports fields, and green dots denote parks. (b) Representative panoramic photo of an Adelaide sports field site. (c) Representative panoramic photo of an Adelaide nature park site

### Vegetation surveys

We surveyed vegetation at all sites between August 14 to 25, 2022, using established methods from White, et al. [24]). In brief, this included assessing 26 points at 1 m intervals across six north-south transects separated by 5 m within each replicate site (6 x 26 = 156 points per site). At each point, we used the plant growth forms “graminoid”, “herb”, “shrub”, and “tree” to record the species richness and proportion of growth forms from ground to canopy, with differentiation at the species level whenever possible. The presence of bare ground (i.e., no vegetation cover including grasses or herbs), litter, and canopy cover were also recorded at each point, due to their potential influence on soil abiotic factors and aerobiome characteristics. The number of points with each of bare ground, litter, and canopy cover were divided by the total points (*n* = 156) to obtain their proportion.

### Rainfall

Rainfall data (in mm) from the recording station in closest proximity to each site was downloaded from the Bureau of Meteorology website [25]. The Bureau of Meteorology utilises > 45 stations across greater metropolitan Adelaide to record daily rainfall. The total rainfall for the seven days prior to the sampling day was aggregated for analysis.

### Air sampling

At each urban greenspace site, air samples were collected over an approximately 8-hour period during the days of May 21-24 and June 11-14, 2022, following the method described in Mhuireach, et al. [11]). We sampled seven sports fields and 14 parks in May, and we sampled four sports fields and eight parks in June. The aerobiome sampling stations were set up on site between 07:00 and 09:00 hours and collected between 15:00 and 17:00.

Each urban greenspace had a single sampling station, a plastic box at a height of 0.3 m from the ground. We opened and placed three sterile, clear plastic Petri dish bases and lids on each station, providing six collection surfaces per site (**Fig. 2**). This method of passive aerobiome sampling has been shown to be as effective as active sampling methods [11]. A field control for each day was generated by holding an additional Petri dish open for 30 seconds at the equipment box. Immediately after the field sampling activity, each Petri dish was sealed, labelled, transported on ice, and then frozen at −20°C until DNA extraction (described below).

**Fig. 2.**
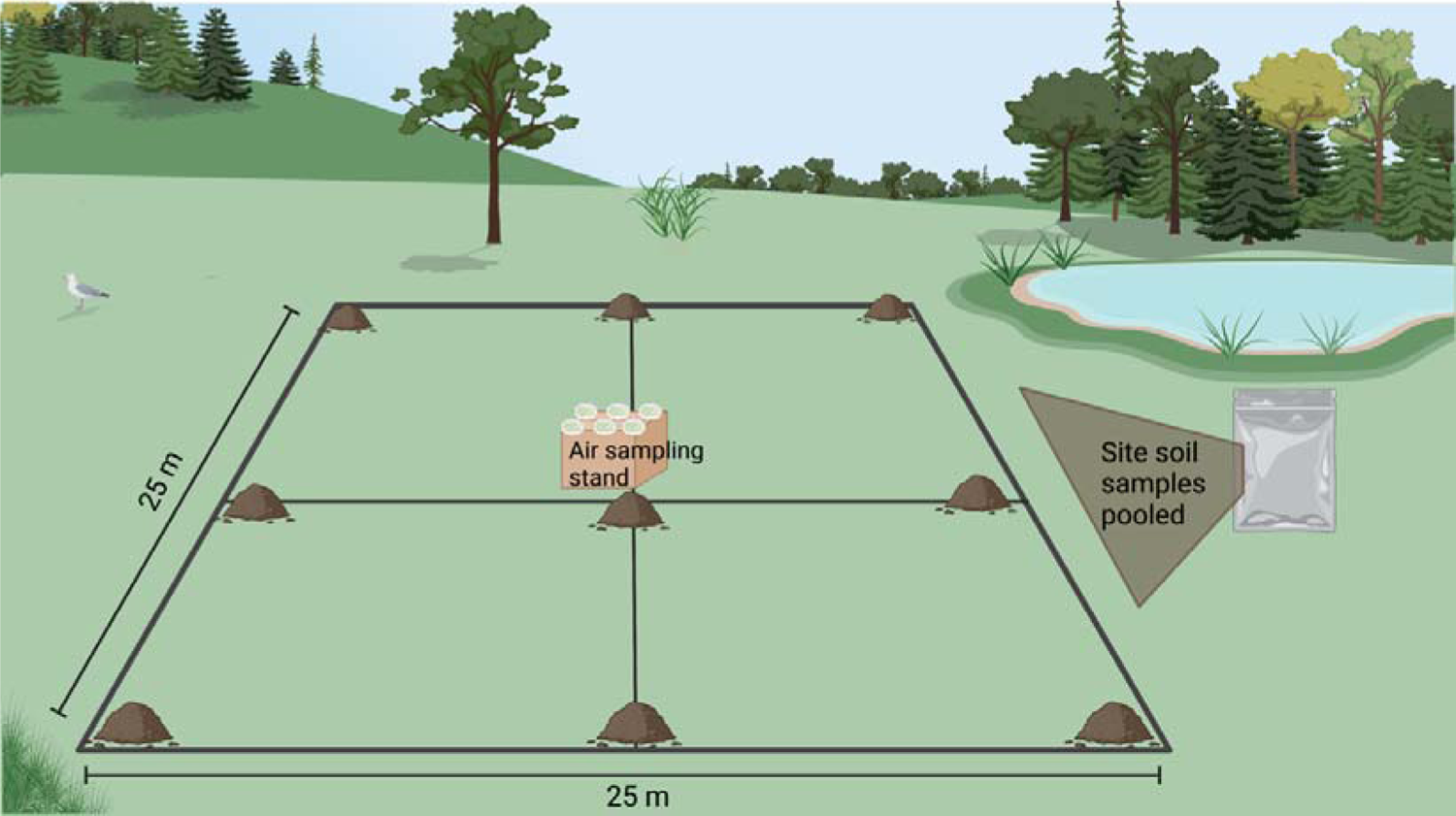
Diagram of sampling area for air and soil sample collection. Soil mounds indicate sample collection points, with all soil samples at one site pooled into a single bag. The air sampling stand with six Petri dish surfaces was placed at the central point. Figure created with Biorender

### Soil sampling

Each 25 x 25 m site area was formatted into nine grid points (**Fig. 2**). We employed Australian Microbiome Initiative [26] sampling protocols. Briefly, this involved using a trowel decontaminated with ethanol and 5% Decon 90 (Decon Laboratories Ltd, Pennsylvania, USA) to collect approximately 50 g of soil from 0-5 cm soil depth at each grid point. The soil samples were then pooled into a sterile plastic bag, homogenised, and the bag sealed. Upon completion of all sampling sites, a minimum of 180 g subsample of each homogenised composite soil sample was placed into new bags and sent to CSBP Soil and Plant Analysis Laboratory (Bibra Lake, Western Australia) for analysis of 23 physical and chemical parameters, including pH, organic carbon, nitrate nitrogen, and cation concentrations (see **Table S1** for all parameter data). Soil moisture (%) was calculated in-house using an oven-drying process as follows: 40 g from each soil sample was transferred to an unsealed metal container, weighed, and placed in an oven at 105°C for 24 hours. Containers with the oven-dried soils were then re-weighed, and the weight lost (= weight of water) as a percentage of total dry mass was calculated.

### eDNA extraction, PCR, sequencing

eDNA extractions and quantifications were performed in the dedicated Biological Sciences laboratory at Flinders University, South Australia. The Petri dishes for each site were opened and swabbed with sterile nylon-flocked swab tips (FLOQSwabs Lot 2011490, Copan Flock Technologies, Bescia, Italy) inside a laminar flow cabinet (Aura PCR PC10000, EuroClone, Milan, Italy). One swab and 40 uL of added sterile phosphate-buffered saline was used for swabbing all six surfaces, except for surfaces that showed visual signs of damage or environmental contamination, for approximately four minutes total using a consistent pattern of swabbing. The tips were cut directly into 2 mL sterilised tubes (QIAGEN, Hilden, Germany). During laboratory preparation for DNA extraction, an extraction blank control was creating using sterile water and processed using the same DNA extraction process as the field site samples.

For air sample DNA extractions, we used the QIAamp DNA Mini Kit (QIAGEN, Hilden, Germany) and followed the manufacturer’s instructions. The extraction concentrations were then quantified using the Quantus fluorometer (Promega, Madison, WI, USA). Samples were sent to South Australia Genomics Centre (SAGC; Adelaide, South Australia) for library preparation via their 16S Metagenomic Sequencing Library Preparation protocol. DNA libraries were QC’d by Tapestation 2200 for size and Qubit for quantity. Equimolar pools were prepared and denatured with a final concentration of 4nM. Pooled libraries were diluted to 8pM (including 10%PhiX) and used for cluster generation. Denaturing and on-board clustering was performed using SAGC’s Illumina protocol v05. Amplification of the bacterial 16S rRNA V3-V4 regions was performed using 314F-806R primers for library preparation that included 25+8 cycles of amplification, and sequencing of amplicon sequence variants was completed on the Illumina Miseq v3 at South Australia Genomics Centre.

### Bioinformatics

From the aerobiome 16s rRNA raw sequence data, amplicon sequence variants (ASVs) were trimmed and filtered using an established Qiime2 pipeline [version 2022.2; 27]. Seven samples had insufficient read counts and were removed from subsequent aerobiome analyses. For the remaining 26 samples, taxonomy was assigned using the onboard Naïve Bayes taxonomic classifier and Greengenes 13-8 database. Sequences were then cleaned using custom code on the R phyloseq package [version 1.42.0; 28] by removing the following: sequences from mitochondria and chloroplasts, taxa that did not occur in at least two samples to reduce artifacts, and ASVs with sums <30. Sequences that were likely of contamination origin were identified and removed using the R decontam package [version 1.18.0; 29] using the function “isNotContaminant” suited for use with low biomass samples.

### Statistical analysis

All statistical analyses were performed using R [version 4.2.3; 30]. To maintain consistency with prior aerobiome studies, statistical significance was set at alpha = 0.05. For each sample we generated statistical data on alpha diversity using Shannon’s Diversity index and Faith’s phylogenetic diversity, beta diversity ordinations using both Aitchison distances and weighted Unifrac, and distance-to-centroid. To prepare phylogenetic trees, sequences were rarefied at the sequencing depth of 12,054, multiple sequence alignment was performed using the program MAFFT [31] with post-cleaning using GBlocks [32], and phylogenetic trees were created using IQTree2 [33] using the “Generalized time-reversible with Gamma rate variation” parameter. Two outlying low values of alpha diversity among sports fields (Kingswood Oval: Shannon index = 2.82, and Daly Oval: Shannon index = 4.64) were omitted from alpha diversity analyses because these outliers were considered unrepresentative at more than two standard deviations from the sports fields group mean. Maps were created using the R ggmap package [version 3.0.2; 34].

To prepare for compositional beta diversity tests, the sequence abundance data was evaluated using R *zCompositions* package [version 1.4.0.1; 35] and zeros were imputed using R *scImpute* package [version 0.0.9; 36]. The resultant abundances were then transformed with centred-log ratio using the R *compositions* package [version 2.0.6; 37], followed by ordination with principal components analysis based on Aitchison distances obtained with the R *vegan* package [version 2.6.4; 38]. In addition, weighted Unifrac distances were calculated with R *vegan* package using the phylogenetic trees described above. Distance-to-centroid analyses were performed using the *betadisper* function in the R *vegan* package. Analysis of Compositions of Microbiomes with Bias Correction using the *ancombc2* function in the R *ANCOMBC* package [version 2.0.3; 39] was performed on untransformed amplicon data for differential abundance analyses. The ANCOMBC algorithm has been shown to minimise bias due to sampling fractions and reduce false discovery rates. Canonical correspondence analyses were performed using the R *CCA* package [version 1.2.1; 40]. The step-based variable selection model was performed using the *ordistep* function in the R *vegan* package. R *ggplot2* package [version 3.4.2; 41] was used for data visualisations.

## RESULTS

We obtained 17.8 million raw reads with 73.6% >Q30 from 33 air samples. After quality control, data from 26 samples with 6,205 bacterial amplicon sequence variants across 33 phyla were utilised in subsequent analyses.

### Aerobiome alpha diversity

Land cover had a weak effect on aerobiome alpha diversity, with sports fields (X = 6.26 ± 0.26) higher than nature parks (X = 5.24 ± 1.27) (Welch Two Sample t-test: *t* = −2.9979, *p* = 0.065; **Fig. 3a, Fig. S1a**). Volume of rainfall in the seven days prior to sampling had a negative effect on the aerobiome alpha diversity (*F* = 5.265, *R^2^* = 0.15, df = 1 and 23, *p* = 0.03; **Fig. 3b, Fig. S1b**). Soil iron negatively influenced the alpha diversity in nature parks (*F* = 22.38, *R^2^* = 0.60, df = 1 and 13, *p* < 0.001; **Fig. 3c, Fig. S1c**), and nature parks had a substantially lower mean concentration of soil iron (X = 60.4 mg/kg ± 39.8) than sports fields (X = 116 mg/kg ± 104). Soil pH had a positive effect on the alpha diversity of both sports fields (*F* = 6.155, *R^2^* = 0.42, df = 1 and 6, *p* = 0.048) and nature parks (*F* = 6.156, *R^2^* = 0.27, df = 1 and 13, *p* = 0.03; **Fig. 3d, Fig. S1d**) (all correlations between soil physicochemical parameters and alpha diversity indices are shown in **Fig. S2 and S3**). Among parks, the diversity of woody plant species had no effect on the aerobiome alpha diversity (*F* = 0.88, *R^2^*= −0.01, df = 1 and 13, *p* = 0.37; **Fig. S4a-b**).

**Fig. 3.**
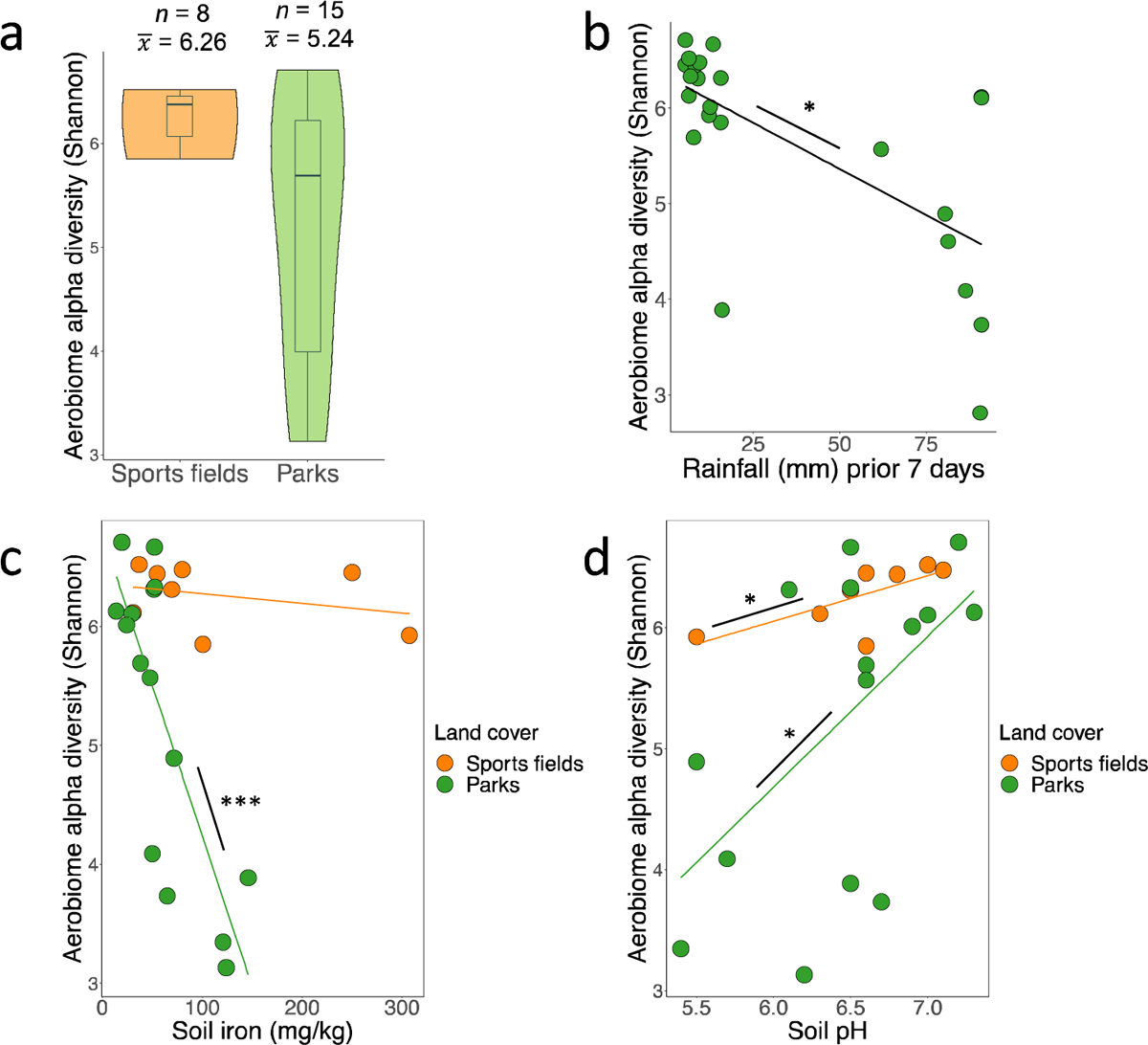
(a) Boxplots of aerobiome alpha diversity by land cover. The y-axis shows the aerobiome alpha diversity calculated by the Shannon diversity index. Boxes show the median and interquartile range, while whiskers extend to the remaining range of data. (b) Relationship of aerobiome bacterial alpha diversity and total rainfall during seven days prior to sampling. (c) Relationship of soil iron with aerobiome alpha diversity. Orange and green lines separately show regressions for sports fields and nature parks, respectively. (d) Relationship of soil pH with aerobiome alpha diversity. Orange and green lines show regressions separately for sports fields and nature parks, respectively

### Aerobiome community composition

Land cover type had a significant influence on the aerobiome community composition. Sports fields had a distinct aerobiome compared to nature parks (PERMANOVA Adonis test: 999 permutations, df = 1, *F* = 3.01, *R*^2^ = 0.12 *p* = 0.001; **Fig. 4a, Fig. S5a**). Sports fields (distance to centroid = 122) and nature parks (distance to centroid = 118.5) had similar levels of homogeneity of aerobiome compositions (ANOVA: *F* = 0.28, df = 1, *p* = 0.60). Rainfall in the week prior to sampling significantly influenced aerobiome community composition across both land cover types (PERMANOVA Adonis test: 999 permutations, df = 1, *F* = 1.84, *R^2^* = 0.07, *p* = 0.016; **Fig. 4b, S5b**). Among nature parks, where the woody plant species diversity (Shannon’s Diversity index) ranged from 0.60 to 1.90, woody plant species diversity had no effect on aerobiome composition (PERMANOVA Adonis test: 999 permutations, df = 1, *F* = 0.80, *R^2^* = 0.05, *p* = 0.82; **Fig. S6a-b**).

**Fig. 4.**
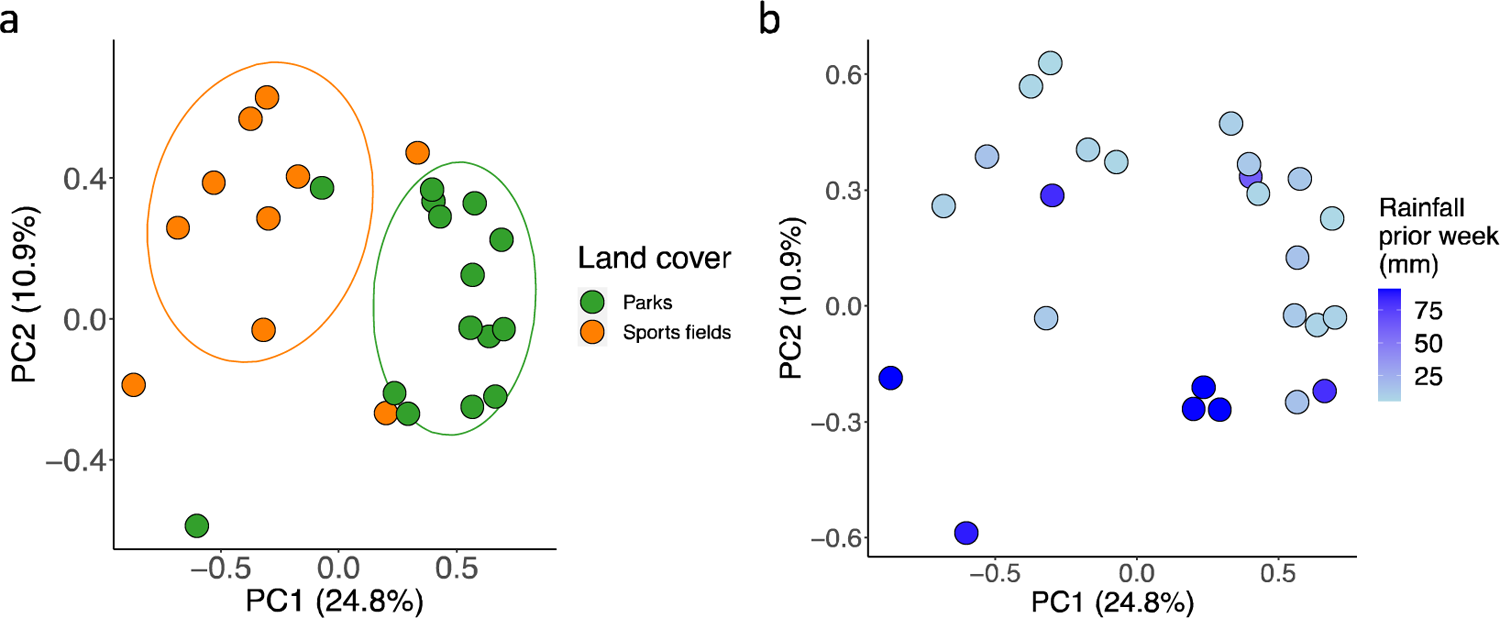
(a) Principal components analysis based on Aitchison distances displaying significant variation in aerobiome community composition between land cover types (PERMANOVA Adonis test: 999 permutations, df = 1, *F* = 3.01, *R*^2^ = 0.12 *p* = 0.001, *n* = 25 sites). (b) Principal components analysis based on Aitchison distances displaying variation in aerobiome community composition by total rainfall volume in the week prior to sampling (PERMANOVA Adonis test: 999 permutations, df = 1, *F* = 1.84, *R^2^* = 0.07, *p* = 0.016, *n* = 25 sites)

The ten phyla with the highest abundances from both land covers showed similar proportions between sports fields and nature parks (**Fig. 5a-b**). Proteobacteria was the dominant phylum in samples from both land covers. Land cover had a strong effect on Desulfobacterota abundances (ANCOM-BC log linear model: log fold change from parks = 2.51, *W* = 3.89, adjusted *p* = 0.002) and a weak effect on Nitrospirota abundances (ANCOM-BC log linear model: log fold change from parks = 2.06, *W* = 2.85, adjusted *p* = 0.052; **Fig. 5c)**, both of which were higher in sports fields aerobiomes. While these phyla were differentially abundant, they were not among the ten phyla with the highest abundances. Among the 353 genera in the dataset, only *Thermomonas* (ANCOM-BC log linear model: log fold change from parks = 2.85, *W* = 3.74, adjusted *p* = 0.03) and an uncultured genus (ANCOM-BC log linear model: log fold change from parks = 3.18, *W* = 3.93, adjusted *p* = 0.03) had significantly greater abundances in sports fields aerobiomes versus nature parks (all genus differential abundances listed **Table S2**).

**Fig. 5.**
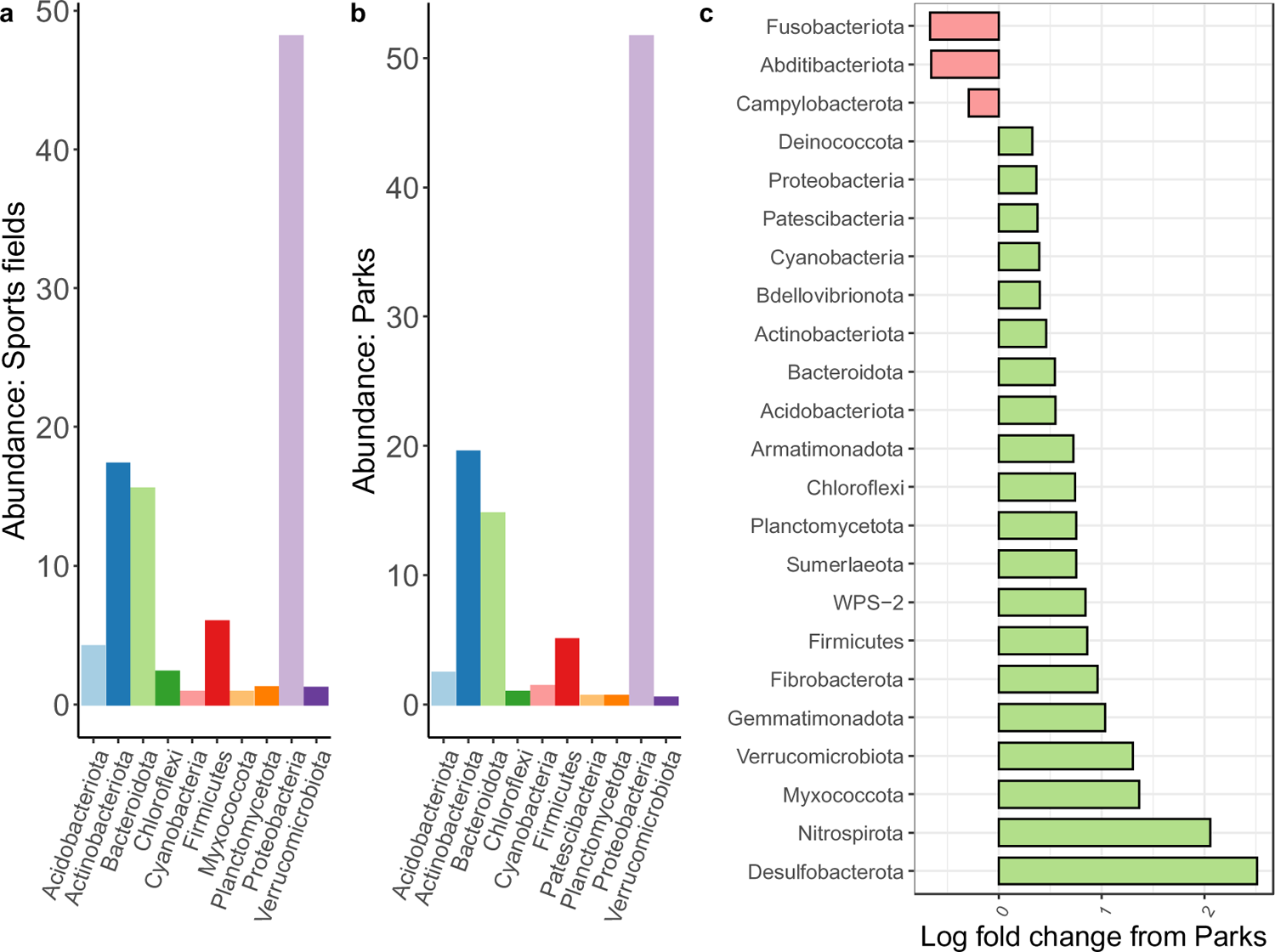
(a) Bar plot showing the phyla with the ten highest mean relative abundances among air samples at sports field sites. Y axis shows the means of the ASV relative abundances from each phylum. (b) Bar plot showing the phyla with the ten highest mean relative abundances among air samples at all park sites. Y axis shows the means of the ASV relative abundances from each phylum. (c) Differential abundance of phyla in aerobiomes by land cover type. The x axis shows the log fold change from parks to sports fields. Red bars indicate a decrease in log fold change, and green bars indicate an increase in log fold change

### Influences of soil physicochemical parameters on aerobiome composition

With data from both land cover types merged, soil physicochemical variables were overall weak predictors of aerobiome composition, and both canonical correspondence analysis (CCA) axes described < 7% (**Fig. 6a**). However, repeating the CCA only in nature parks (*n* = 15) showed that soil iron had a strong effect on the aerobiome composition (ANOVA: *F* = 1.308, df = 1, *p* = 0.04; **Fig. 6b**). Among sports fields (*n* = 10), soil physicochemical variables showed no significant effect on aerobiome composition among sports fields (**Fig. 6c**).

**Fig. 6.**
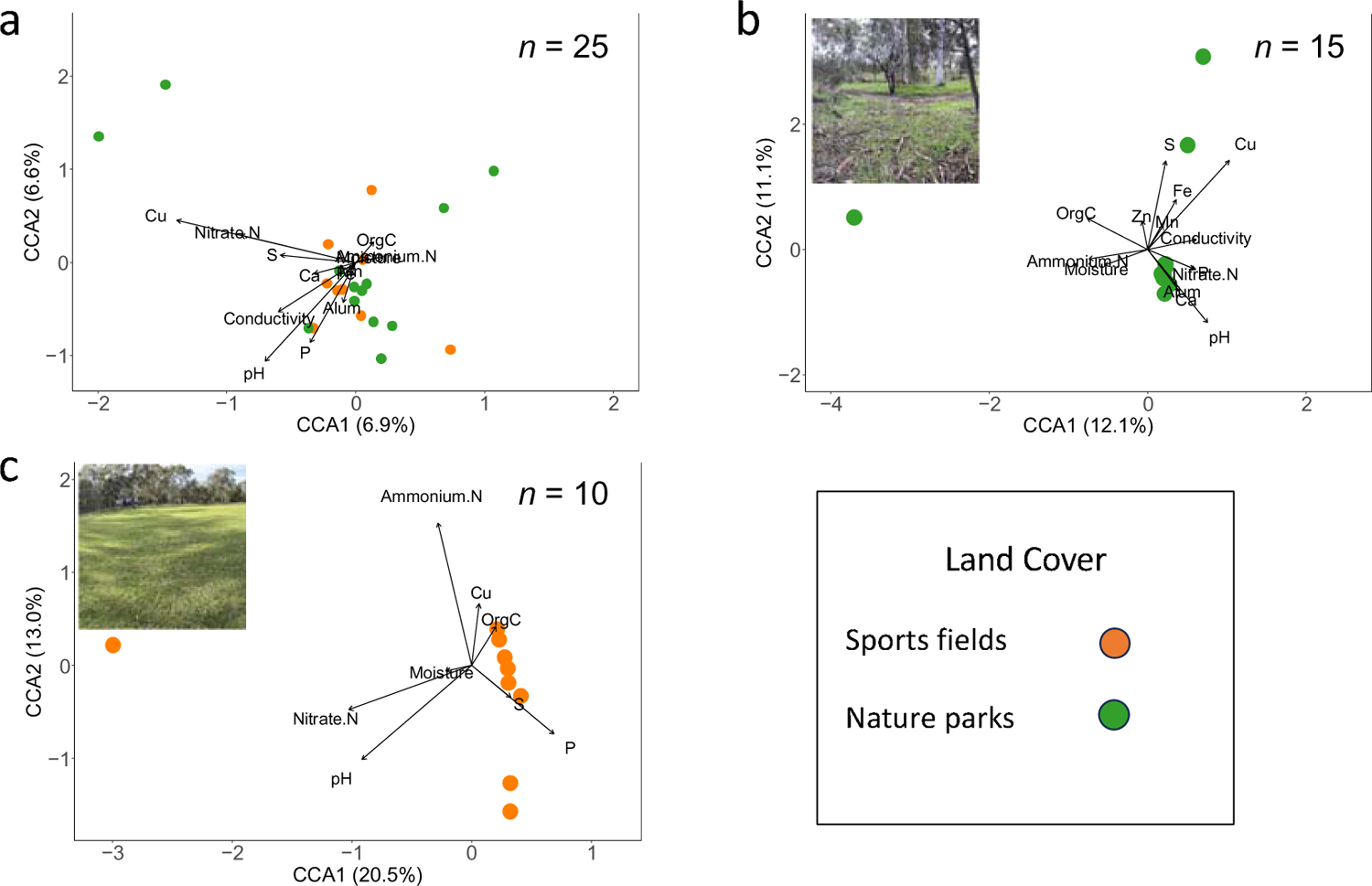
(a) Canonical correspondence analysis (CCA) of aerobiome community composition constrained by soil physicochemical variables across all sites. (b) CCA of aerobiome community composition of nature parks constrained by soil physicochemical variables. (c) CCA of aerobiome community composition of sports fields constrained by soil physicochemical variables. In (a-c), arrow lengths indicate strength of constraint by each explanatory variable, and arrowheads show the direction of increase. Orange points indicate sports fields and green points indicate nature parks; *n* is the number of sites

## DISCUSSION

Urban greenspaces provide city residents with opportunities to gain access to nature and exposure to potentially health-supporting biodiverse airborne bacterial communities (aerobiomes). Therefore, describing the characteristics of aerobiomes in urban greenspaces, and identifying the ecological factors that influence them, can support greenspace management and future planning. Here, we found that near-surface aerobiomes of sports fields had distinctly different bacterial communities than nature parks. However, levels of bacterial diversity were influenced more by soil physicochemical characteristics (primarily soil pH and iron) and rainfall than by the land cover type and the woody vegetation diversity. The vegetation and soils of urban greenspaces, especially sports fields, appear to be managed in such a way that results in consistent characteristics of the airborne microbial communities. Because such characteristics are important for urban greenspace development, we examined the influence of four key ecological variables on urban greenspace aerobiomes, each of which is discussed below.

### Effects of soil physicochemical parameters on aerobiomes

We show that soil physicochemical parameters had a greater influence on the alpha diversity of aerobiomes than on the community composition. Though iron played the strongest role among the parameters in shaping the aerobiome community, its effects were moderate and were limited to nature parks. On the other hand, soil pH and iron concentrations had strong effects on the aerobiome alpha diversity. Previous research has demonstrated that abiotic features (e.g., pH) are important drivers in shaping microbial community composition in soils, sometimes taking precedence over plant-based factors [42]. The importance of soil abiotic factors in shaping bacterial alpha diversity is recognised in soils, but to our knowledge has not previously been translated to aerobiomes. Although here we did not analyse soil microbiomes, the bacteria in soils are well-known contributors to aerobiomes [17, 18]. Our results suggest that maintaining soil pH close to neutral (e.g., via liming) and lowering (or otherwise counteracting) elevated soil iron levels could increase the aerobiome alpha diversity. Notably, pH associated with aerobiome alpha diversity regardless of the land cover. Thus, human exposure to salutogenic microbial diversity could potentially be modulated through landscape maintenance strategies that focus on soil pH across both sports fields and nature parks.

It is important to note that manually altering soil pH could affect wider ecosystem functions. For instance, landscape maintenance efforts to substantially shift soil pH could affect the growth and distribution of plant species in the ecosystem [43] and alter soil carbon cycling [44] potentially impacting the entire food web. Furthermore, exposure to greater alpha diversity might not invariably produce salutogenic effects; the community composition and their associated functional traits may also play a role. For instance, species richness (and evenness) might be 100-fold higher in one sample compared to another but, in theory, could contain 100-fold more pathogens with high relative abundances. In such a scenario, alpha diversity will be considerably higher, but so will the sample’s pathogenic potential. Nonetheless, altering soil properties to enhance the aerobiome in highly-managed anthropogenic environments, such as sports fields, could contribute to human immunoregulation, whilst minimising adverse ecological impacts.

### Effects of land cover type on aerobiomes

We found that greenspace land cover weakly influenced aerobiome alpha diversity, with intensely-managed grassy sports fields having higher alpha diversity than less-managed nature parks. While related aerobiome studies are limited, our findings are consistent with Delgadol Baquerizo, et al. [16] who observed higher soil bacterial alpha diversity in urban greenspaces with more intense land management than in adjacent, more natural sites. It may be possible that sports field management techniques, such as regular fertilisation, introduce soil conditions that select for a wider range of bacterial inhabitants. Because soil bacterial communities are shaped by a complex network of biotic and abiotic factors, future research can examine the unique properties of sports field management that may influence the soil and airborne bacterial communities.

Furthermore, we found that the aerobiome community composition of sports fields was distinct from nature parks. Each of the top ten most abundant aerobiome phyla had higher abundances in sports fields than in nature parks. The greatest abundance difference was observed in the phylum Desulfobacterota, which includes taxonomic classes with a preference for anaerobic environments [45] and the capacity to use sulphur and iron as terminal electron acceptors. Sports field soils had almost double the concentration of both sulphur and iron than nature parks. As such, sports fields soil conditions may be selecting for bacteria such as Desulfobacterota with the capacity to utilise sulphur and iron in metabolic processes.

### Effects of woody plant species diversity on aerobiomes

Previous studies have shown that wooded urban habitats with more complex vegetation had higher aerobiome alpha diversity than grassy habitats [12, 13]. Leaf surfaces can offer key sources of airborne bacteria [46], and greater vegetation complexity can provide additional leaf surface. Among our nature park sites, however, the diversity of woody plant species did not influence the aerobiome alpha diversity. The seasonal conditions prior to sampling (May-June 2022) with late autumn to winter rainfall may have been a factor in ‘washing off’ and reducing the contribution of phyllosphere microbiota to aerobiomes.

In addition, our finding that soil abiotic characteristics had a stronger effect than nearby vegetation could have been influenced by our sampling height. While we sampled at 0.3 m height, Mhuireach, et al. [12], who found that vegetation affected aerobiomes, sampled at 2 m height. Aerobiomes, including pathogenic taxa, are known to stratify vertically, with greater alpha diversity closer to the ground [13, 20]. It is, therefore, possible that vegetation may have a greater effect on aerobiomes above 0.3 m. Future studies should evaluate how ecological features, including detailed soil and vegetation properties, impact aerobiomes at varying heights.

### Effects of rainfall on aerobiomes

We observed that increased rainfall in the week prior to sampling associated with a reduction in aerobiome alpha diversity. Rainfall is known to reduce the volume of bioaerosols [47, 48]. Whether bioaerosol volume reduction links with a decrease in bacterial diversity likely depends on factors such as the typical aerosol particle size present to which bacteria can attach, which can influence which specific taxa may be removed by rainfall [49].

Simultaneously, rainfall can disperse surface soil bacteria into the air through the impact of raindrops on soils [50]. When previously airborne bacteria are removed by rain, and soil-borne bacteria are dispersed into the air, the aerobiome can experience a shift in composition [49]. Indeed, we observed that increased rainfall associated with a difference in aerobiome composition. Future studies should employ repeated sampling methods at urban greenspaces to gain further insight into specific compositional changes associated with rainfall, and whether those changes could be beneficial or harmful to human health.

### Limitations

Our study examined the bacteria only among air samples. Because soils make key contributions to adjacent aerobiomes, future studies that directly compare aerobiome and soil microbiome compositions could provide further insights into quantity and mechanistic theory of bacterial transfer, and subsequent human exposure, in urban greenspaces. Although we focused on bacteria, future assessment of fungi among urban greenspace aerobiome datasets could also give further insights into the contributions of greenspace exposure to human health and disease. In addition, wind can carry distant airshed bacteria and have a substantial influence on the aerobiome; further studies could examine the effects of wind on urban greenspace aerobiomes to which humans may be exposed.

## Conclusions

We performed a novel analysis of airborne microbial communities across two types of urban greenspaces: sports fields and nature parks. We show that the near-surface aerobiomes at 0.3 m sampling height were influenced by land cover type, recent rainfall, and soil physicochemical characteristics (i.e., soil pH and iron) but not by woody plant species diversity. We also show that soil pH strongly affects the bacterial diversity in the air regardless of greenspace land cover type. Greenspaces are an integral part of urban design, providing opportunities for city residents to gain greater exposure to natural biodiversity.

Urban greenspace planners and restoration ecologists rely on data to manage ecological variables such as woody plant diversity and soil physicochemical parameters. Our findings should further assist the development of greenspace design initiatives that aim to harness the health-promoting effects of biodiverse microbial exposures.

## DECLARATIONS

### Funding

This work was supported by funding from the Flinders Foundation, Australian Research Council [grants LP190100051] and the New Zealand Ministry of Business Innovation and Employment [grant UOWX2101].

### Competing interests

The authors have no competing interests to declare that are relevant to the content of this article.

### Data availability

The datasets generated during and/or analysed in the current study are available in Supplementary information, and all datasets and custom R code are available on figshare at 10.6084/m9.figshare.24112917.

### Authors’ contributions

All authors contributed to the study conception and design. Material preparation, data collection and analysis were performed by Joel Brame, Craig Liddicoat, and Martin Breed. The first draft of the manuscript was written by Joel Brame and all authors commented on further versions of the manuscript. All authors read and approved the final manuscript.

## Notes

http://doi.org/10.6084/m9.figshare.24112917

### Competing Interest Statement

The authors have declared no competing interest.

## REFERENCES

[1] J. Robinson, J. Mills, and M. Breed, “Walking ecosystems in microbiome-inspired green infrastructure: an ecological perspective on enhancing personal and planetary health,” Challenges, vol. 9, no. 2, p. 40, 2018, 10.3390/challe9020040.

[2] G. Rook, “Regulation of the immune system by biodiversity from the natural environment: an ecosystem service essential to health,” Proceedings of the National Academy of Sciences, vol. 110, no. 46, pp. 18360–18367, 2013, 10.1073/pnas.1313731110.

[3] M. Jimenez, et al., “Associations between nature exposure and health: A review of the evidence,” International Journal of Environmental Research and Public Health, vol. 18, no. 9, p. 4790, 2021.

[4] L. Von Hertzen, I. Hanski, and T. Haahtela, “Natural immunity: biodiversity loss and inflammatory diseases are two global megatrends that might be related,” EMBO Reports, vol. 12, no. 11, pp. 1089–1093, 2011, 10.1038/embor.2011.195.

[5] B. Wintle et al., “Global synthesis of conservation studies reveals the importance of small habitat patches for biodiversity,” Proceedings of the National Academy of Sciences, vol. 116, no. 3, pp. 909–914, 2019.

[6] M. Kondo, J. Fluehr, T. McKeon, and C. Branas, “Urban green space and its impact on human health,” International Journal of Environmental Research and Public Health, vol. 15, no. 3, p. 445, 2018.

[7] M. Marselle, S. Lindley, P. Cook, and A. Bonn, “Biodiversity and health in the urban environment,” Current Environmental Health Reports, vol. 8, no. 2, pp. 146–156, 2021.

[8] T. Haahtela, “A biodiversity hypothesis,” Allergy, vol. 74, no. 8, pp. 1445–1456, 2019, 10.1111/all.13763.

[9] M. Roslund et al., “A Placebo-controlled double-blinded test of the biodiversity hypothesis of immune-mediated diseases: Environmental microbial diversity elicits changes in cytokines and increase in T regulatory cells in young children,” Ecotoxicology and Environmental Safety, vol. 242, p. 113900, 2022.

[10] T. Alfven, C. Braun-Fahrlander, B. Brunekreef, E. von Mutius, J. Riedler, and A. Scheynius, “Allergic diseases and atopic sensitisation in children related to farming and anthroposophic lifestyle - the PARSIFAL study,” (in English), *Allergy*, Correction vol. 62, no. 4, pp. 455–455, Apr 2007, 10.1111/j.1398-9995.2007.01361.x.

[11] G. Mhuireach et al., “Urban greenness influences airborne bacterial community composition,” Sci Total Environ, vol. 571, pp. 680–7, Nov 15 2016, 10.1016/j.scitotenv.2016.07.037.

[12] G. Mhuireach, H. Wilson, and B. Johnson, “Urban aerobiomes are influenced by season, vegetation, and individual site characteristics,” EcoHealth, vol. 18, pp. 331–344, 2021.

[13] J. Robinson et al., “Exposure to airborne bacteria depends upon vertical stratification and vegetation complexity,” Scientific Reports, vol. 11, no. 1, p. 9516, 2021.

[14] R. Bowers, S. McLetchie, R. Knight, and N. Fierer, “Spatial variability in airborne bacterial communities across land-use types and their relationship to the bacterial communities of potential source environments,” The ISME journal, vol. 5, no. 4, pp. 601–612, 2011.

[15] Z. Baruch et al., “Characterising the soil fungal microbiome in metropolitan green spaces across a vegetation biodiversity gradient,” Fungal Ecology, vol. 47, p. 100939, 2020.

[16] M. DelgadolBaquerizo et al., “Ecological drivers of soil microbial diversity and soil biological networks in the Southern Hemisphere,” Ecology, vol. 99, no. 3, pp. 583–596, 2018.

[17] J. Robinson and M. Breed, “The aerobiome–health axis: a paradigm shift in bioaerosol thinking,” Trends Microbiol., 2023.

[18] E. Brodie, T. DeSantis, J. Parker, I. Zubietta, Y. Piceno, and G. Andersen, “Urban aerosols harbor diverse and dynamic bacterial populations,” Proceedings of the National Academy of Sciences, vol. 104, no. 1, pp. 299–304, 2007.

[19] A. Woo et al., “Temporal variation in airborne microbial populations and microbially-derived allergens in a tropical urban landscape,” Atmospheric Environment, vol. 74, pp. 291–300, 2013.

[20] J. Robinson, C. Cando-Dumancela, C. Liddicoat, P. Weinstein, R. Cameron, and M. F. Breed, “Vertical stratification in urban green space aerobiomes,” Environmental Health Perspectives, vol. 128, no. 11, p. 117008, 2020.

[21] E. Franchitti, C. Caredda, E. Anedda, and D. Traversi, “Urban Aerobiome and Effects on Human Health: A Systematic Review and Missing Evidence,” Atmosphere, vol. 13, no. 7, p. 1148, 2022.

[22] B. Tripathi, J. Stegen, M. Kim, K. Dong, J. Adams, and Y. Lee, “Soil pH mediates the balance between stochastic and deterministic assembly of bacteria,” The ISME Journal, vol. 12, no. 4, pp. 1072–1083, 2018.

[23] J. Mills et al., “Revegetation of urban green space rewilds soil microbiotas with implications for human health and urban design,” Restoration Ecology, vol. 28, pp. S322-S334, 2020.

[24] A. White et al., “AUSPLOTS Rangelands Survey Protocols Manual,” 2012.

[25] Bureau of Meteorology. “Climate Data Online.” http://www.bom.gov.au/climate/data/ (accessed 2022).

[26] A. Bissett, et al., “Introducing BASE: the Biomes of Australian Soil Environments soil microbial diversity database,” GigaScience, vol. 5, no. 1, pp. s13742-016-0126-5, 2016.

[27] E. Bolyen et al., “Reproducible, interactive, scalable and extensible microbiome data science using QIIME 2,” Nat. Biotechnol., vol. 37, no. 8, pp. 852–857, 2019.

[28] P. McMurdie and S. Holmes, “phyloseq: an R package for reproducible interactive analysis and graphics of microbiome census data,” PloS One, vol. 8, no. 4, p. e61217, 2013.

[29] N. Davis, D. Proctor, S. Holmes, D. Relman, and B. Callahan, “Simple statistical identification and removal of contaminant sequences in marker-gene and metagenomics data,” Microbiome, vol. 6, pp. 1–14, 2018.

30. *R: A language and environment for statistical computing.* (2023). R Foundation for Statistical Computing, Vienna, Austria.

[31] K. Katoh and D. Standley, “MAFFT multiple sequence alignment software version 7: improvements in performance and usability,” Molecular Biology and Evolution, vol. 30, no. 4, pp. 772–780, 2013.

[32] G. Talavera and J. Castresana, “Improvement of phylogenies after removing divergent and ambiguously aligned blocks from protein sequence alignments,” Systematic Biology, vol. 56, no. 4, pp. 564–577, 2007.

[33] L. Nguyen, H. Schmidt, A. Von Haeseler, and B. Minh, “IQ-TREE: a fast and effective stochastic algorithm for estimating maximum-likelihood phylogenies,” Molecular Biology and Evolution, vol. 32, no. 1, pp. 268–274, 2015.

[34] D. Kahle and H. Wickham, “ggmap: spatial visualization with ggplot2,” R J., vol. 5, no. 1, p. 144, 2013.

[35] J. Palarea-Albaladejo and J. Martín-Fernández, “zCompositions—R package for multivariate imputation of left-censored data under a compositional approach,” Chemometrics and Intelligent Laboratory Systems, vol. 143, pp. 85–96, 2015.

[36] W. Li and J. Li, “An accurate and robust imputation method scImpute for single-cell RNA-seq data,” Nat. Commun., vol. 9, no. 1, p. 997, 2018.

[37] K. Van den Boogaart and R. Tolosana-Delgado, ““Compositions”: a unified R package to analyze compositional data,” Computers & Geosciences, vol. 34, no. 4, pp. 320–338, 2008.

[38] J. Oksanen et al., “vegan: Community Ecology Package. R package version 2.5-7. 2020,” ed, 2022.

[39] H. Lin and S. Peddada, “Analysis of compositions of microbiomes with bias correction,” Nat. Commun., vol. 11, no. 1, p. 3514, 2020.

[40] I. González, S. Déjean, P. G. Martin, and A. Baccini, “CCA: An R package to extend canonical correlation analysis,” Journal of Statistical Software, vol. 23, pp. 1–14, 2008.

41. *Ggplot2: Elegant graphics for data analysis (2nd ed)*. (2016). Springer-Verlag, New York.

[42] N. Fierer, “Embracing the unknown: disentangling the complexities of the soil microbiome,” Nat. Rev. Microbiol., vol. 15, no. 10, pp. 579–590, 2017.

[43] F. Seaton et al., “Fifty years of reduction in sulphur deposition drives recovery in soil pH and plant communities,” Journal of Ecology, vol. 111, no. 2, pp. 464–478, 2023.

[44] A. Malik et al., “Land use driven change in soil pH affects microbial carbon cycling processes,” Nat. Commun., vol. 9, no. 1, p. 3591, 2018.

[45] C. Murphy et al., “Genomic characterization of three novel Desulfobacterota classes expand the metabolic and phylogenetic diversity of the phylum,” Environmental Microbiology, vol. 23, no. 8, pp. 4326-4343, 2021.

[46] R. Bowers, A. Sullivan, E. Costello, J. Collett Jr, R. Knight, and N. Fierer, “Sources of bacteria in outdoor air across cities in the midwestern United States,” Applied and Environmental Microbiology, vol. 77, no. 18, pp. 6350–6356, 2011.

[47] Y. Li, R. Lu, W. Li, Z. Xie, and Y. Song, “Concentrations and size distributions of viable bioaerosols under various weather conditions in a typical semi-arid city of Northwest China,” Journal of Aerosol Science, vol. 106, pp. 83–92, 2017.

[48] V. Després, et al., “Primary biological aerosol particles in the atmosphere: a review,” Tellus B: Chemical and Physical Meteorology, vol. 64, no. 1, p. 15598, 2012.

[49] G. Jang, C. Hwang, and B. Cho, “Effects of heavy rainfall on the composition of airborne bacterial communities,” Frontiers of Environmental Science & Engineering, vol. 12, pp. 1–10, 2018.

[50] Y. Joung, Z. Ge, and C. Buie, “Bioaerosol generation by raindrops on soil,” Nat. Commun., vol. 8, no. 1, p. 14668, 2017.

