## Supplementary Information for "Urban greenspace aerobiomes are shaped by soil conditions and land cover type"

**Table S1**: Sample data of 23 soil physicochemical variables (separate file on Figshare, doi: 10.6084/m9.figshare.24112917)

**Table S2**: Genus-level bacterial differential abundances between sports fields and nature parks (separate file on Figshare, doi: 10.6084/m9.figshare.24112917)

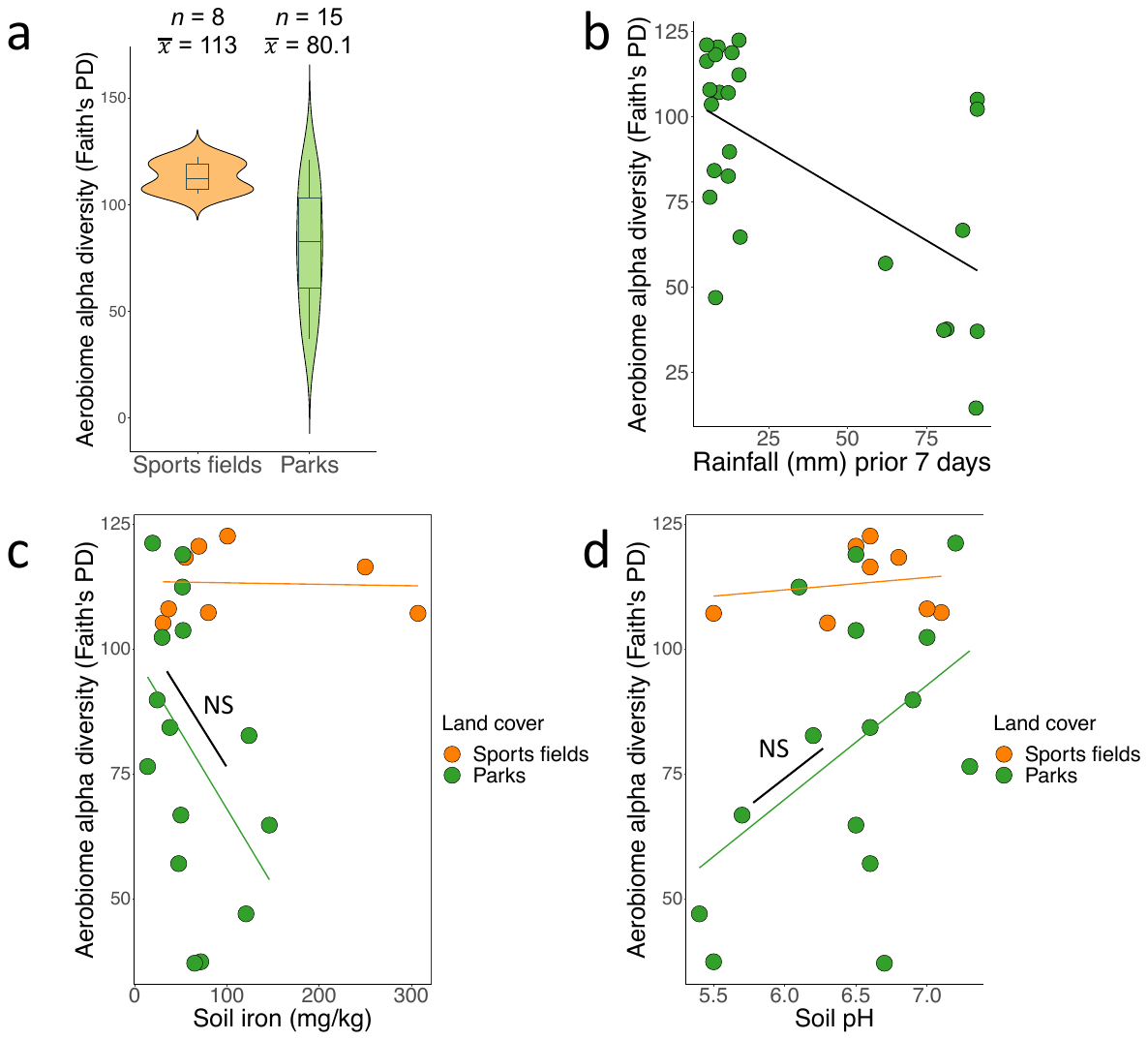

**Fig. S1**. (a) Boxplots of aerobiome alpha diversity by land cover. The y-axis shows the aerobiome alpha diversity calculated by Faith’s phylogenetic diversity (PD). Boxes show the median and interquartile range, while whiskers extend to the remaining range of data. (b) Relationship of aerobiome bacterial alpha diversity (Faith’s PD) and total rainfall volume during seven days prior to sampling. (c) Relationship of soil iron with aerobiome alpha diversity (Faith’s PD). Orange and green lines separately show regressions for sports fields and nature parks, respectively. (d) Relationship of soil pH with aerobiome alpha diversity (Faith’s PD). Orange and green lines show regressions separately for sports fields and nature parks, respectively

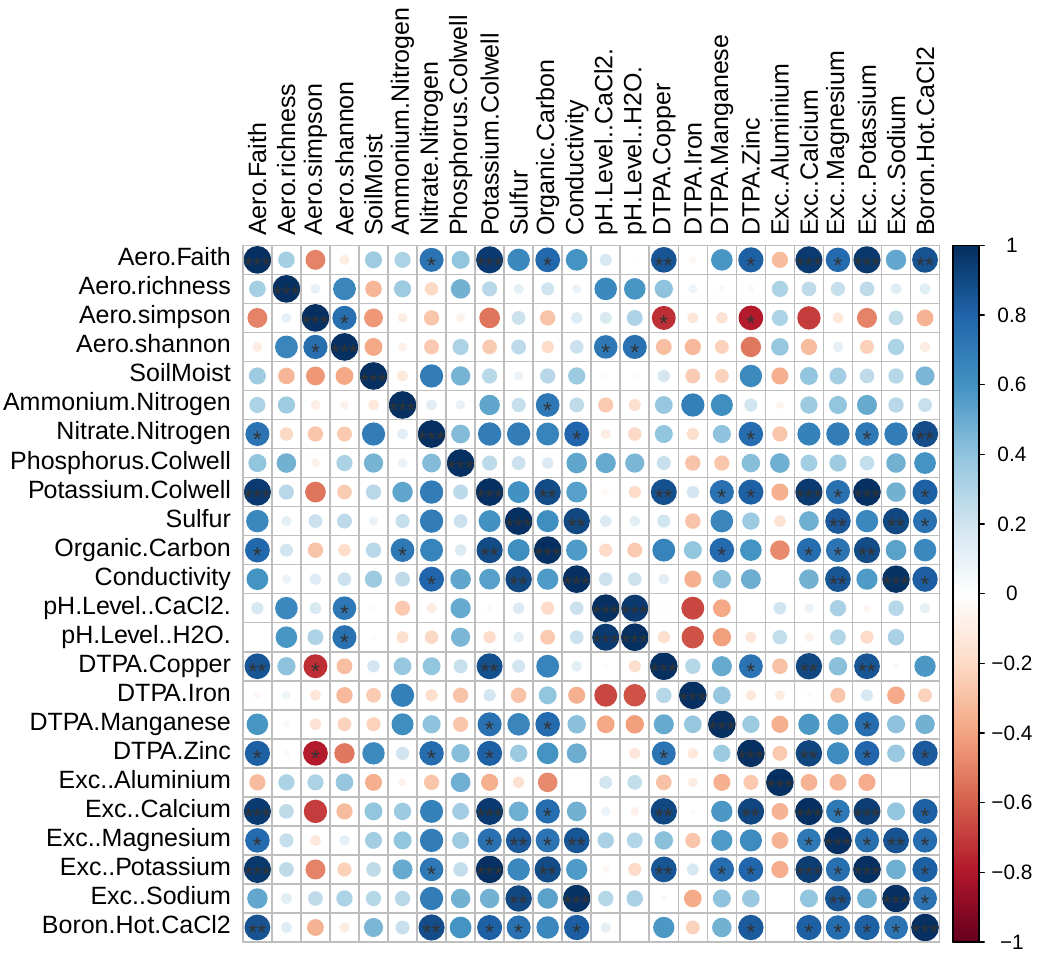

**Fig. S2**. Pearson correlation coefficients between four aerobiome alpha diversity indices and soil physicochemical parameters in sports fields. * indicates significance at *p* < 0.05, and ** indicates significance at *p* < 0.01.

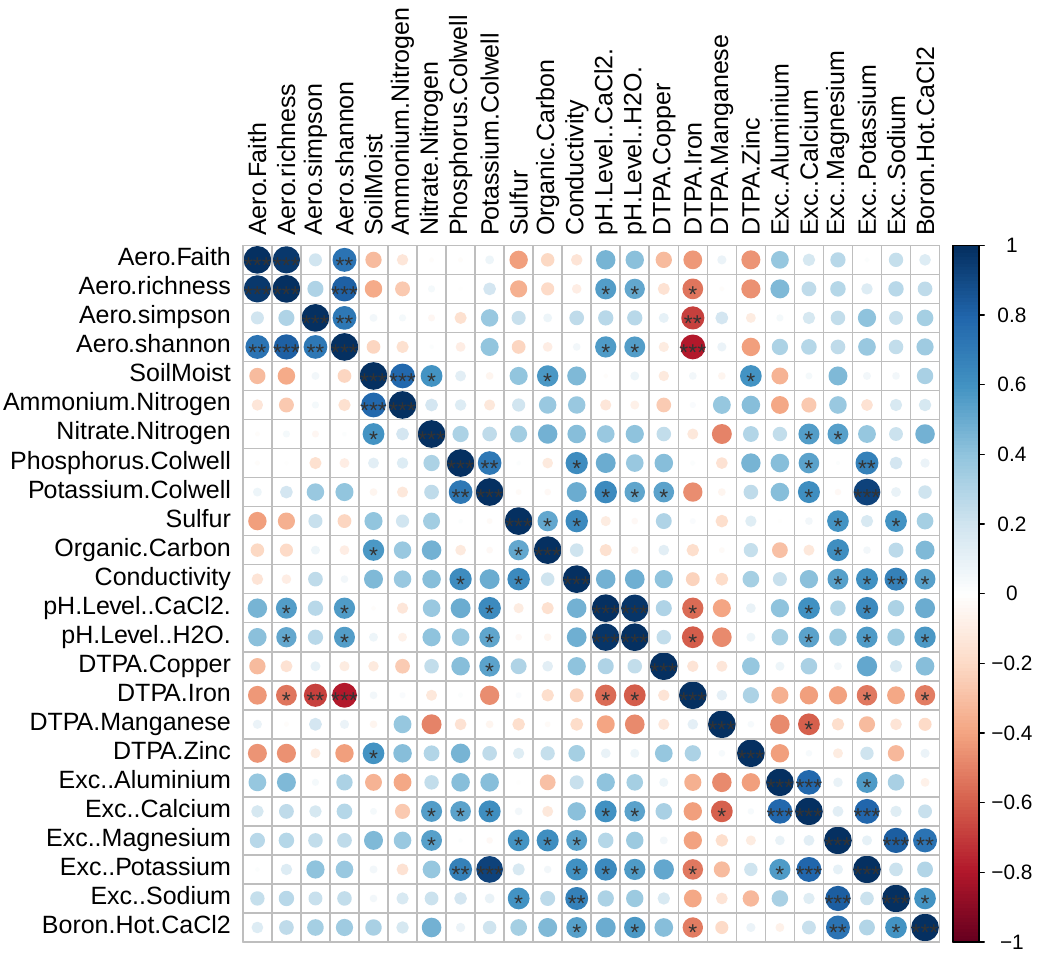
**Fig. S3**. Pearson correlation coefficients between four aerobiome alpha diversity indices and soil physicochemical parameters in nature parks. * indicates significance at *p* < 0.05, ** indicates significance at *p* < 0.01, and *** indicates significance at *p* < 0.001

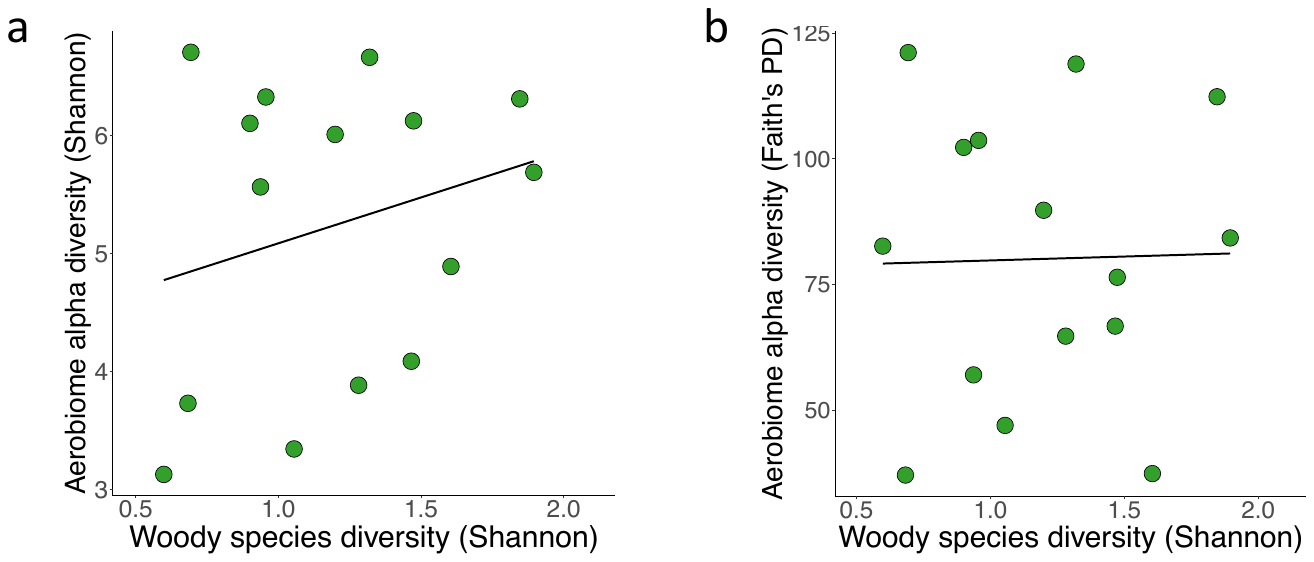

**Fig. S4**. (a) Relationship of woody plant species diversity (Shannon) with aerobiome alpha diversity using Shannon index. (b) Relationship of woody plant species diversity (Shannon) with aerobiome alpha diversity using Faith’s phylogenetic diversity.

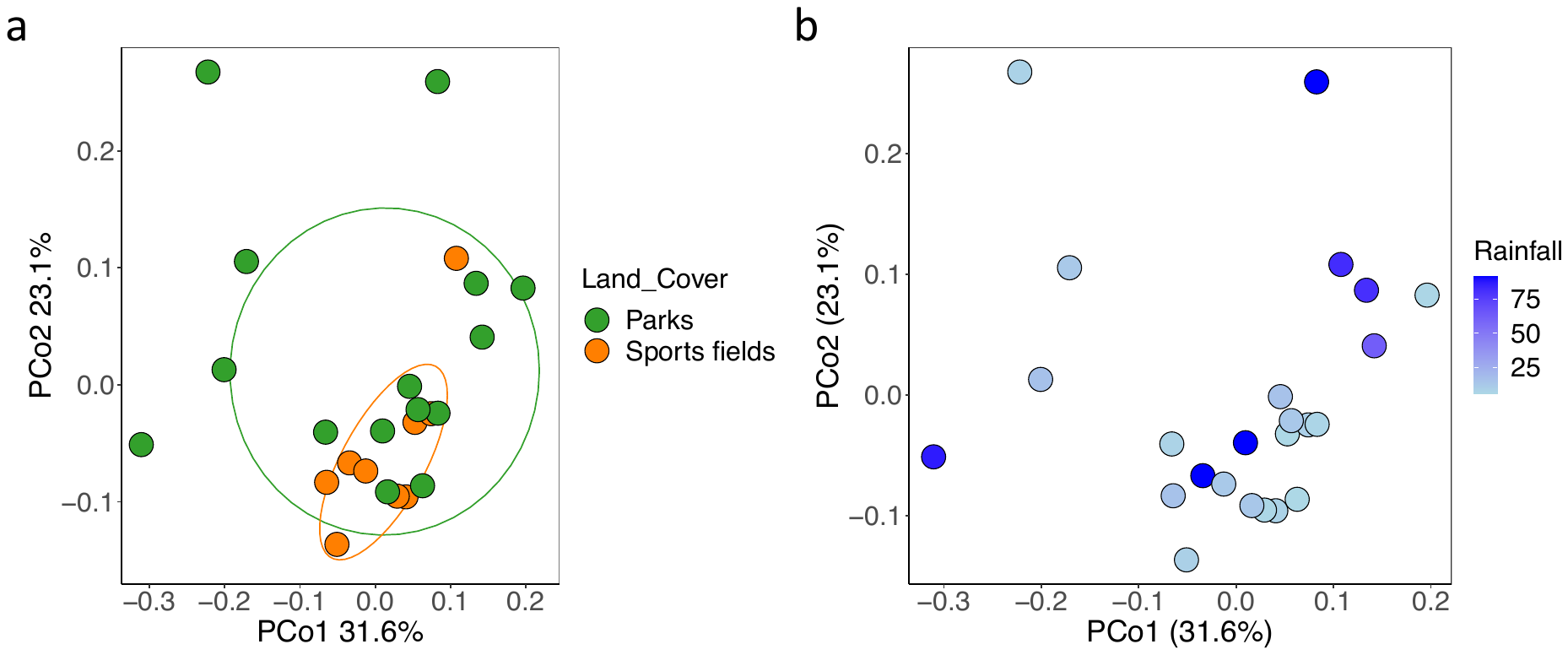

**Fig. S5**. (a) Principal coordinates analysis based on weighted Unifrac distances displaying variation in aerobiome community composition between land cover types (PERMANOVA Adonis test: 999 permutations, df = 1, *F* = 1.423, *R*^2^ = 0.06 *p* = 0.168, *n* = 25 sites). (b) Principal coordinates analysis based on weighted Unifrac distances displaying variation in aerobiome community composition by total rainfall volume in the week prior to sampling (PERMANOVA Adonis test: 999 permutations, df = 1, *F* = 1.524, *R*^2^ = 0.06 *p* = 0.12, *n* = 25 sites).

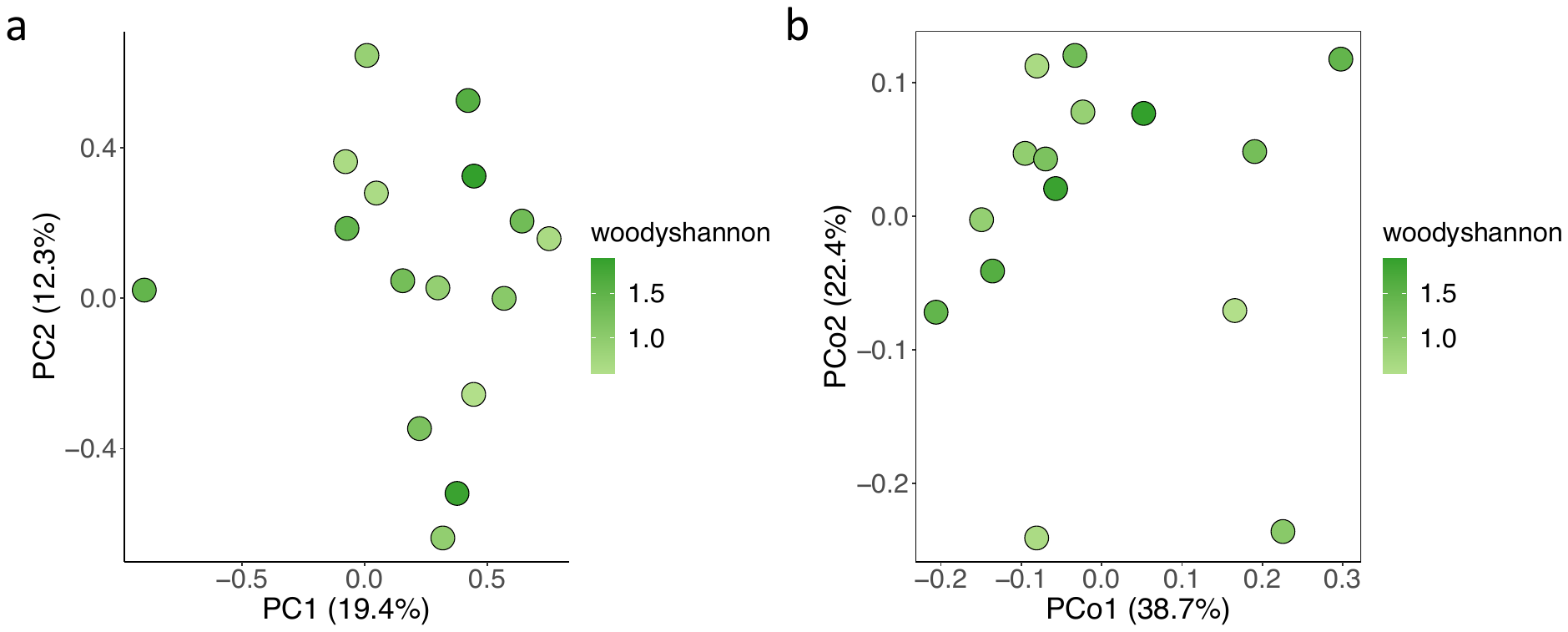

**Fig. S6**. (a) Principal components analysis based on Aitchison distances displaying variation in aerobiome community composition of parks with varying Shannon diversity of woody plant species (PERMANOVA df = 1, *F* = 0.80, *R^2^* = 0.05, *p* = 0.82, *n* = 14 sites). (b) Principal coordinates analysis based on weighted Unifrac distances displaying variation in aerobiome community composition of parks with varying Shannon diversity of woody plant species (PERMANOVA df = 1, *F* = 0.50, *R^2^* = 0.04, *p* = 0.86, *n* = 14 sites)
